# Definition of regulatory elements and transcription factors controlling immune cell gene expression at single cell resolution using single nucleus ATAC-seq

**DOI:** 10.1101/2024.01.09.574769

**Authors:** Pengxin Yang, Ryan Corbett, Lance Daharsh, Juber Herrera Uribe, Kristen A. Byrne, Crystal L. Loving, Christopher Tuggle

## Abstract

The transcriptome of porcine peripheral blood mononuclear cells (PBMC) at single cell (sc) resolution is well described, but little is understood about the cis-regulatory mechanism behind scPBMC gene expression. Here, we profiled the open chromatin landscape of porcine PBMC using single nucleus ATAC sequencing (snATAC-seq). Approximately 22% of the identified peaks overlapped with annotated transcription start sites (TSS). Using clustering based on open chromatin pattern similarity, we demonstrate that cell type annotations using snATAC-seq are highly concordant to that reported for sc RNA sequencing (scRNA-seq). The differentially accessible peaks (DAPs) for each cell type were characterized and the pattern of accessibility of the DAPs near cell type markers across cell types was similar to that of the average gene expression level of corresponding marker genes. Additionally, we found that peaks identified in snATAC-seq have the potential power to predict the cell type specific transcription starting site (TSS). We identified both transcription factors (TFs) whose binding motif were enriched in cell type DAPs of multiple cell types and cell type specific TFs by conducting transcription factor binding motif (TFBM) analysis. Furthermore, we identified the putative enhancer or promoter regions bound by TFs for each differentially expressed gene (DEG) having a DAP that overlapped with its TSS by generating cis-co-accessibility networks (CCAN). To predict the regulators of such DEGs, TFBM analysis was performed for each CCAN. The regulator TF-target DEG pair predicted in this way was largely consistent with the results reported in the ENCODE Transcription Factor Targets Dataset (TFTD). This snATAC-seq approach provides insights into the chromatin accessibility landscape of porcine PBMCs and enables discovery of TFs predicted to control DEG through binding regulatory elements whose chromatin accessibility correlates with the DEG promoter region.

## 1. INTRODUCTION

The pig is of great economic importance since it is a crucial source of protein and meat world-wide. Pigs are also a valuable model for translational biomedical research resulting from their high similarity to human in size, genomics, immunology and physiology (Groenen et al., 2012; Dawson et al., 2013; Lunney et al., 2021). Benchmark epigenetic studies analyzing multiple tissues have characterized porcine cis-regulatory elements (Kern et al., 2021; Zhao et al 2021), and cis-regulatory elements were reported to have higher conservation with human than between human and mouse (Zhao et al., 2021). Peripheral blood mononuclear cells (PBMCs) are an extensively studied sample in -omics and biomedicine since they are easy to collect and express numerous functional markers (Vandiedonck, 2018). In addition, PBMCs include many of the major cells in porcine immunity. Thus, PBMCs can serve as a great resource to monitor individual immune homeostasis.

Transcriptomes of porcine PBMCs at both bulk and single cell resolution using RNA-seq and single-cell RNA sequencing (scRNA-seq), respectively, have been reported (Herrera-Uribe et al., 2021). But gene expression alone provides limited information in terms of gene regulation. The chromatin accessibility of human hematopoietic cells was profiled by applying transposase-accessible chromatin sequencing in single nuclei (snATAC-seq) (Buenrostro et al., 2018). But such single cell/nuclei epigenomic landscapes in porcine PBMCs has not been reported. Genome-wide chromatin accessibility can reflect not only the transcription factor (TF) binding but also the regulatory capacity at the open chromatin region (Klemm et al. 2019). snATAC-seq allows inference of gene expression for genes with low RNA abundance that are hard to detect by scRNA-seq methods. snATAC-seq also makes it possible to predict future transcription since the openness of chromatin likely happens prior to any transcription. It has been reported that snATAC-seq has comparable ability to scRNA-seq in terms of cell type annotation and may be able to detect more distinct cell types compared to scRNA-seq (Miao et al. 2021). Moreover, by calculating co-accessibility, snATAC can predict long-range chromatin interaction which is unique compared to scRNA (Pliner et al. 2018). Therefore, elucidating the chromatin accessibility of porcine PBMCs can provide necessary information to identify the cell type specific cis-regulatory elements that, through chromatin interactions, have the capacity to regulate transcription. Such identified regulomes can enhance the understanding of the epigenetic mechanisms governing the establishment of cell differentiation and cell functionality.

Here, to elucidate the genome-wide epigenetic landscape of porcine PBMCs and identify the cis-regulatory mechanism governing the known cell type specific gene expression of porcine peripheral immune cells, we profiled the chromatin accessibility of porcine PBMCs by applying snATAC-seq. Cell types were annotated by manual gene marker-based annotation using snATAC alone or by integration with our published PBMC scRNAseq data (Herrera-Uribe et al., 2021). Differentially accessible peaks (DAPs) for genes in each annotated cell type were identified, and TFs enriched in annotated cell type DAPs were predicted. Cis-co-accessibility networks (CCANs) were generated to predict the long-range chromatin interaction regulating nearby genes, and TF binding motif (TFBM) enrichment analysis on the DAPs in each CCAN was performed.

## 2. RESULTS

### 2.1 Single-cell chromatin landscape of healthy porcine immune cells

The chromatin accessibility landscape of PBMCs collected from two healthy 6-month male pigs was profiled by performing a microfluidics-based snATAC-seq via 10x Genomics Chromium platform. We generated 4 snATAC-seq libraries from two replicate samples per pig and sequenced them via Illumina Novaseq 6000 sequencing runs. We generated chromatin accessibility profiles from 20,861 nuclei.

The expected fragment size distribution periodicity and TSS enrichment in each of the four datasets were identified (Supplementary Fig 1A). The average median TSS enrichment score across the four snATAC-seq datasets was 17.99 (16.56-19.20) (Supplementary Fig 1B-E). Nuclei doublets from each dataset were detected and filtered out (1,466 doublets) using ArchR (Supplementary Fig 2A-L) (Granja et al. 2021). A set of 110,444 high quality peaks with an average length of 1,086 bp were identified and used for quantifying Tn5 transposase cut sites for each dataset. 78.3% of snATAC peaks (86,494) overlapped with peaks derived from ATAC-seq of bulk sorted porcine PBMC populations (Corbett et al., manuscript in prep), demonstrating high concordance of chromatin accessibility between ATAC-seq on a bulk population and snATAC-seq (Supplementary Fig 3A). To further characterize identified peaks, the proportion of detected peaks in each genomic region was calculated (Fig 1A). Briefly, 22% of peaks were within 3kb of the TSS of a gene. The Tn5 insertion frequency was re-quantified for each of the snATAC-seq datasets to create a feature matrix and Seurat object. To check consistency across the four datasets, we randomly selected one region and noted that the peaks across different datasets were highly uniform with each other (Supplementary Fig 4 A-D). Then low-quality nuclei were filtered out based on total number of fragments in the nuclei, number of peaks in the nuclei, nucleosome signal and TSS enrichment score for each dataset. The four datasets were then merged and integrated to generate an evenly distributed snATAC dataset comprising 17,207 nuclei (Supplementary Fig 5A-B). The 2nd to 30th latent semantic indexing (LSI) component was used for clustering analysis since the first LSI component is highly correlated with sequencing depth (Supplementary Fig 5C)(Stuart et al., 2021). Consequently, 35 clusters of nuclei, with at least 1,444 differentially accessible peaks (DAPs) in all pairwise clusters, were identified using a shared nearest neighbor clustering algorithm (Seurat/Signac) and visualized on Uniform Manifold Approximation and Projection (UMAP) (Fig 1B, Supplementary Table 1-2). Overall, there was no obvious dataset-specific clusters though there were a few clusters (cluster 10, 21, 23, 25, 28 and 34) mainly composed of nuclei from either 6798.2x or 6800.2x dataset likely due to the fact that the number of nuclei in those two datasets were approximately four times that of the other two datasets (Supplementary Fig 5D-H). This demonstrated that the batch effect was effectively removed.

**Figure 1.**
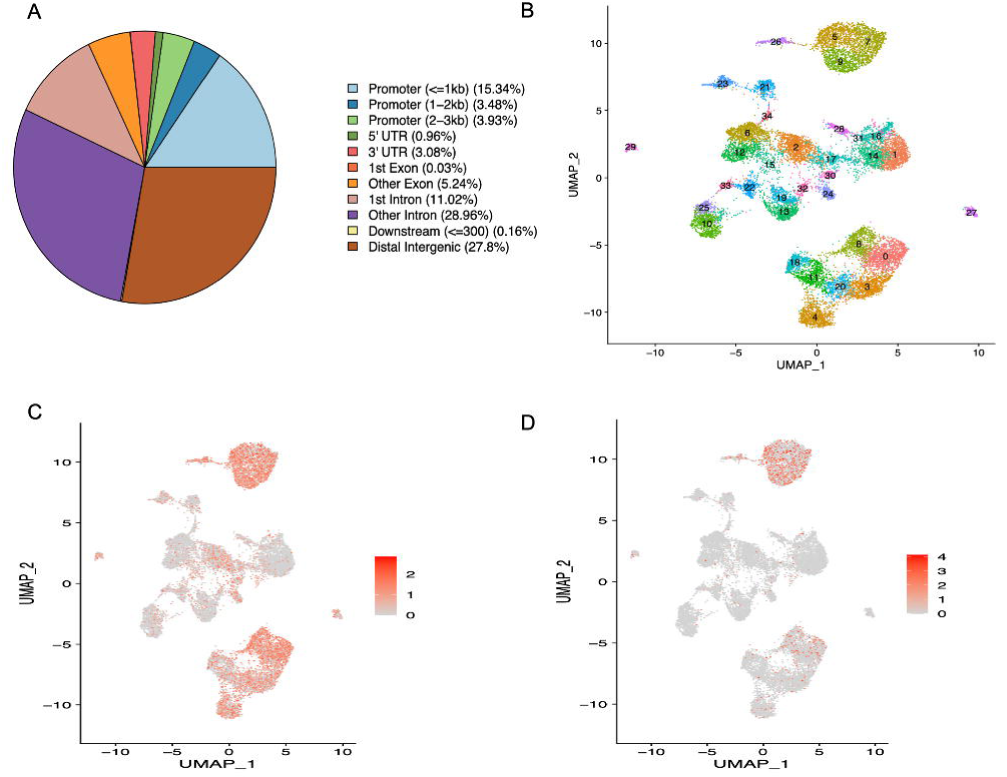
Major porcine peripheral blood mononuclear cell types identified through single nucleus chromatin accessibility profiles. A: Pie chart showing the proportion of indicated genomic regions detected as peaks in snATAC-seq dataset B. UMAP plot of 17,207 nuclei isolated from PBMC subjected to snATAC-seq and separated into 35 clusters based on similarity of the chromatin accessibility pattern. Each point represents a single nucleus. C: UMAP plot of estimated *CD86* gene activity by counting the Tn5 transposase cutting sites in all fragments near *CD86* gene (< 2000bp of TSS). D: UMAP plot of the chromatin accessibility at a cluster differentially accessible peak (DAP) (13-138488083-138490791) whose nearest gene is monocyte cell marker *CD86*.

### 2.2 Cell type annotation of porcine PBMC using chromatin accessibility landscape

The cell type of each cluster was manually classified by estimating the gene activity from the number of Tn5 transposase cutting sites within the identified peaks of +/-2kb of TSS of well-defined canonical marker genes. A series of marker genes (Herrera-Uribe et al., 2021) for specific cell types were investigated for their overall gene activity in each cluster (see example of *CD86*, a myeloid cell marker, in Fig 1C, other marker gene activity patterns are shown in Supplementary Figs 6-11). We also identified differentially accessible peaks (DAPs) for each cluster (average log2FC>0.25, p_val_adj < 0.05) and approximately 18% (20,070 of 110,444) of the unique cis-elements were found to be differentially accessible in at least one cluster (Supplementary Table 3 DE.pig.resol2.4.2.30.findallmarker.onlypos.p0.05.txt). These DAPs whose nearest gene was a known cell-type-marker gene used in Herrera-Uribe et al., 2021 were annotated and used to estimate gene activity (Fig 1D, Supplementary Fig 12, Supplementary Table 4,). (see “Method” section). A cell was predicted to express a cell type functional gene if it demonstrated measurable gene activity. Consequently, clusters were assigned into 12 cell types (Fig 2A-2B). Seven clusters (0, 3, 4, 8, 11, 18, 20) were identified as B cells, one cluster (27) as antibody-secreting cells (ASCs), three clusters as monocytes (5, 7, 9), one cluster (29) as plasmacytoid dendritic cells (pDCs), one cluster (26) as conventional dendritic cells (cDCs), three clusters (2, 6, 12) as CD4+ αβ T cells (CD4posab), six clusters (13, 19, 23, 24, 30, 32) as CD8αβ+ αβ T cells (CD8abPOSab), one cluster (10) as natural killer cells (NK), three clusters (17, 21, 34) as T cells, two clusters (25, 33) as a mixture of CD8abPOSab and NK cells (CD8abPOSabT_NK), four clusters (1, 14, 16, 28) as CD2-γδ T cells (CD2negGD), and one cluster (22) as CD2+ γδ T-cells (CD2posGD). The latter two cell types (CD2-γδ T cells and CD2+ γδ T-cells) were annotated based on a collection of marker genes (see “Methods”). Finally, two clusters (15, 31) were classified as unknown cells and no further analysis completed.

**Figure 2.**
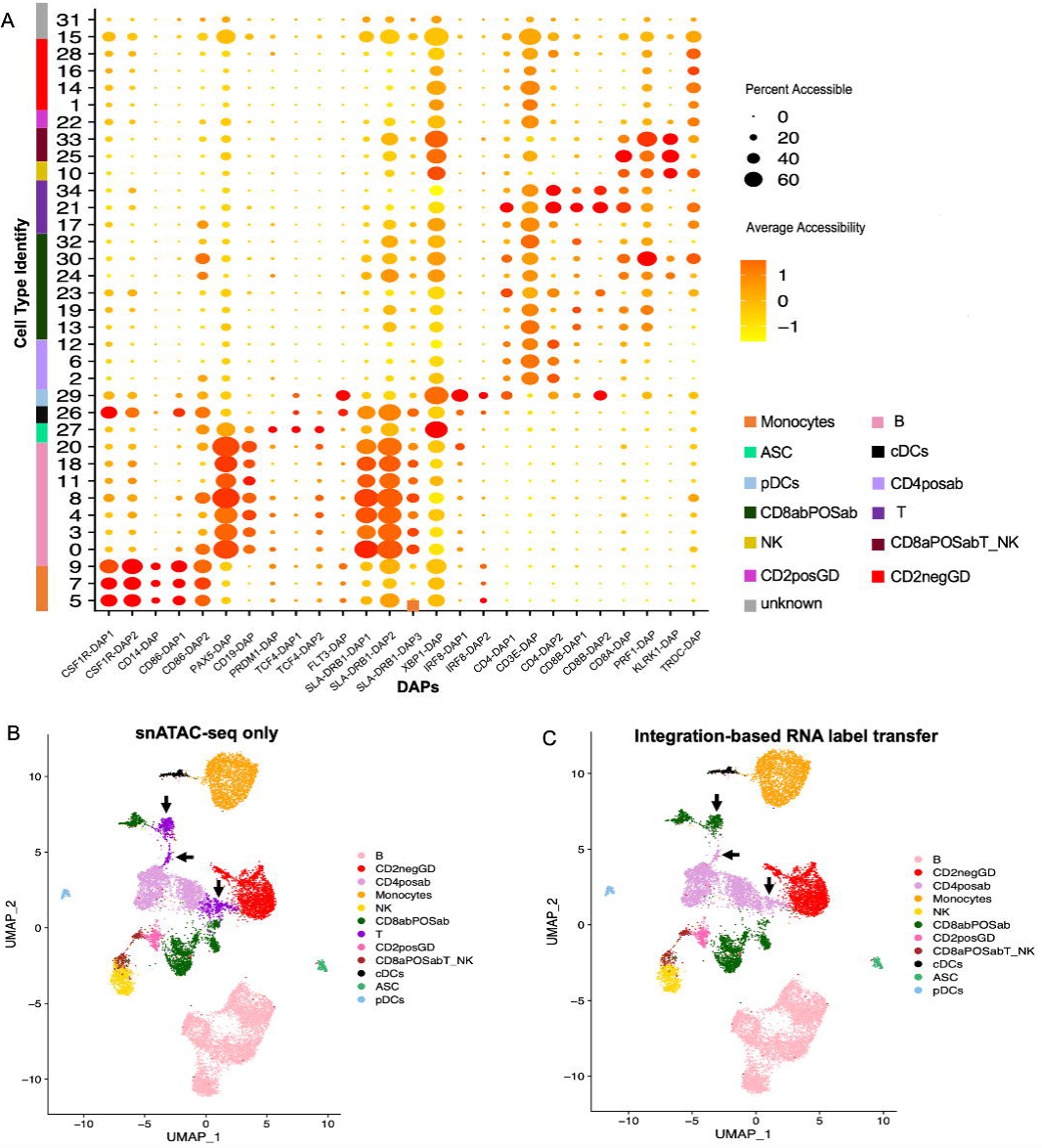
Cluster annotations delineated through integration snATAC-seq estimated gene activity and scRNA-seq gene expression was highly concordant with annotations derived from snATAC-seq data alone. A: Dotplot visualizing the chromatin accessibility at 26 cluster differentially accessible peaks (DAP) near canonical genes indicative of cell type in the 35 clusters derived from snATAC-seq (Fig 1B). The genomic coordinates of the DAP genes on x-axis are listed in Supplementary Table 6. The size of the dot represents the fraction of nuclei having chromatin accessibility at the matching DAP on x-axis in each cluster. The larger dot indicates a higher percentage of nuclei with the region accessible in respective cluster. The color of the dot denotes the average chromatin accessibility level across all nuclei in the respective cluster (red is high). B: UMAP plot of the snATAC-seq dataset with cell types annotated from Fig 2A. Estimated gene activity score was calculated using two criteria described in materials and methods and cells in clusters were annotated into 13 cell types: B cells, CD2-γδ T-cells (CD2negGD), CD4+ αβ T cells (CD4posab), Monocytes, natural killer cells (NK), CD8αβ+ αβ T cells (CD8abPOSab), T cells (T), CD2+ γδ T-cells (CD2posGD), CD8abPOSa/NK cells (CD8abPOSabT_NK), conventional dendritic cells (cDCs), antibody secreting cells (ASCs), plasmacytoid dendritic cells (pDCs) and unknown cells (not shown). C: UMAP plot of the snATAC-seq dataset labeled with cell types predicted by integrating gene activity scores from snATAC-seq dataset (Fig 2A) and previously published porcine PBMC expression levels from scRNA-seq dataset. Clusters were classified into 12 cell types: B cells, CD2negGD, CD4posab, Monocytes, NK, CD8abPOSab, CD2posGD, CD8abPOSabT_NK, cDCs, ASC, pDCs and unknown cells (not shown).

Interestingly, leveraging some cluster-specific cis-elements near known gene markers provided equivalent or even better cell type classification than using only overall gene activity (Fig 1C-1D, Supplementary Fig 12, see “Methods”). For instance, cluster 0, 3, 4, 8, 11, 18 and 20, which were annotated as B cells, demonstrated relatively high *CD19* gene activity measured using all cis-elements nearby yet almost exclusive chromatin accessibility at a cis-element near this known B cell marker *CD19* (Supplementary Fig 12A-B). Similarly, the chromatin accessibility of a potential cis-element near monocyte marker *CSF1R* in monocyte clusters 5, 7 and 9 was more unique to monocytes than the estimated *CSF1R* gene activity (Supplementary Fig 12C-12D). Likewise, a possible cis-element near *CD3E*, a known T cell marker, was more specific to all 16 T cells clusters compared to non-T cell clusters than the overall *CD3E* gene activity (Supplementary Fig 12E-12F). Finally, there is a cis-element near the *CD4* gene in CD4+ αβ T cells, which enabled annotation of cluster 2, 6 and 12 as CD4+ αβ T cells more confidently and specifically than utilizing the overall gene activity of *CD4* (Supplementary Fig 12G-12H).

### 2.3 Integration of porcine PBMC snATAC dataset with comparable scRNA dataset

To further validate the cell type annotation and/or possibly revise it, the estimated gene activity from the snATAC dataset was integrated with our previously published scRNA-seq dataset (Supplementary Fig 13A) from porcine PBMCs (Herrera-Uribe et al., 2021) by identifying cross-modality pairwise “anchors” between the two datasets and transferring the scRNAseq annotation label (Stuart et al. 2019) to the snATACseq clusters. Most of the clusters (26 out of 35 clusters) had a median prediction score over 70% (Supplementary Fig 14, Supplementary Table 5 celltype.prediction.score.table). For further functional comparisons, 1,164 nuclei with a prediction score lower than 0.5 were filtered out from the snATAC-seq dataset, leaving 16,043 nuclei (93%) for final annotation and characterization.

The resulting cluster annotations assigned using gene expression from scRNAseq dataset and estimated gene activity of canonical genes for different immune cell types (Fig 2B) was almost identical to the predicted cell type labels in the scRNA dataset (Fig 2C) with the exception of clusters 17, 21 and 34. We decided to annotate the snATAC-seq clusters based on the cell type predicted after integration with scRNA-seq for the following reasons. First, the cell types predicted for cluster 17, 21 and 34 using the integrated analysis (Fig 2C) were more specific than cell types annotated using only cluster DAPs near known cell type markers (Fig 2B). Second, we compared chromatin accessibility of clusters annotated based on predicted cell types (Fig 2C) in peaks shared with those obtained from bulk ATAC-seq of bulk-sorted porcine PBMCs using principal component analysis and found they demonstrated high consistency with each other (Supplementary Fig 3B-C). In addition, an erythrocyte cluster was identified, but this will not be further discussed since there was only one cell predicted to be in this group. Finally, the integration did not resolve the two clusters annotated as a mixture of CD8αβ+ αβ T/NK, nor provide further annotation of the two unknown clusters. Overall, the annotation obtained from integrating snATAC-seq with scRNA-seq clustering of matched cell populations across datasets provided additional biological support for assignment of the snATACseq clusters using only snATACseq data.

### 2.4 Characterization of cell type specific cis-regulatory elements

To identify the cell type specific regulatory genomic regions, DAPs more accessible in specific cell types were detected by performing a Wilcoxon Rank Sum test in Seurat (logfc > 0.25, p_val_adj < 0.05). Consequently, we identified 11,872 unique cell type-specific DAPs across 11 cell types (Table 1; Supplementary Table 7 celltype.DAP.summary) and these DAPs were significantly enriched for DAPs in comparable bulk-sorted porcine PBMC cell populations (Supplementary Fig 3D). Such identified DAPs can be used to identify cis regulatory elements that are associated with specific cell type expression patterns and potentially contribute to the differential expression (DE) of nearby genes in the respective cell. We found that the cell type predicted by the accessibility pattern of identified cis-elements near marker genes in each cell type (Fig 3A) was similar to that predicted by the gene expression pattern of matching marker genes (Fig 3B). For example, we identified a cis-element region that overlaps the TSS of *CSF1R*, a monocyte marker gene, that is significantly more accessible in monocytes which potentially govern DE of *CSF1R* in monocyte cells in scRNA-seq dataset (1st column in Fig 3B). The fact that this cis-element is also accessible in the cDCs might explain the moderate expression of *CSF1R* in cDCs cells (1st column in Fig 3B). Interestingly, when this *CSF1R* DAP is plotted based on the frequency of Tn5 insertion events, this DAP is in the middle of *CSF1R* (Fig 3C). There are five *CSF1R* transcripts sharing three TSSs identified in pigs on Ensembl (Fig 3D), and two of them (for *CSF1R*-201 and *CSF1R*-202) have TSSs overlapping with this *CSF1R* DAP. Then, we extended the evaluation to all DAPs overlapping with a TSS of gene cell markers in Herrera-Uribe et al., 2021. Similarly, we identified only TSS of *CD8A*-201 was within the DAP whose nearest gene is *CD8A* among three transcripts of *CD8A* (Supplementary Fig 15B).

**Figure 3.**
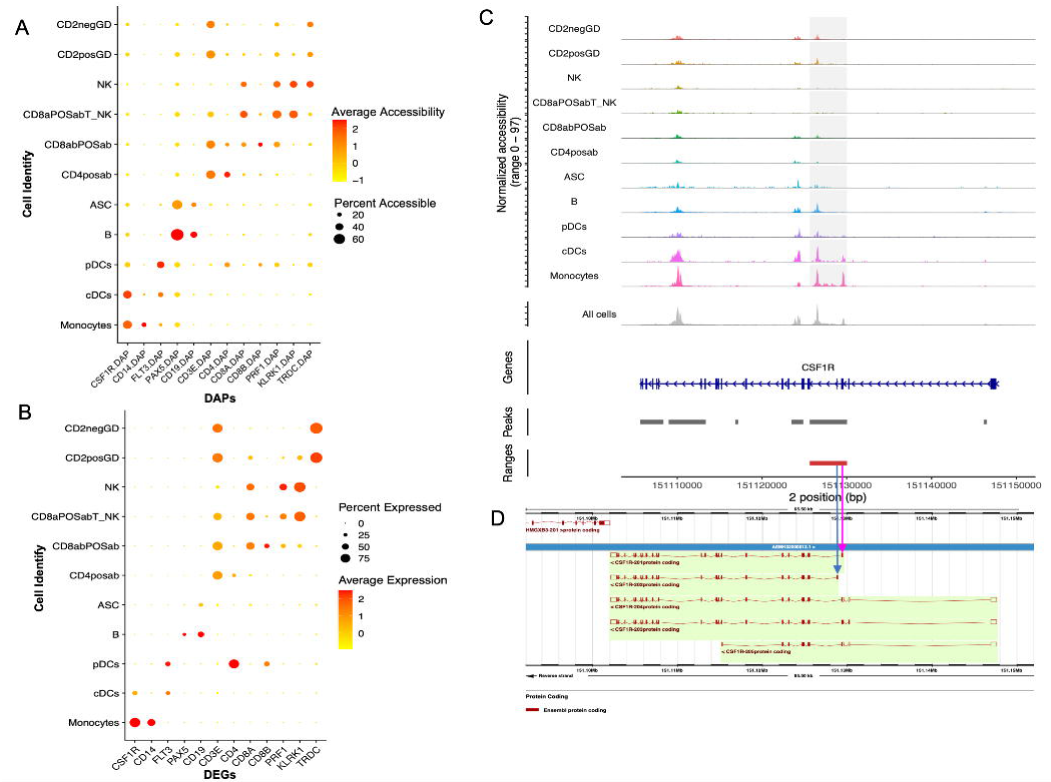
Commensurate patterns of differentially accessible peaks and expression of nearby genes in porcine PBMC. A: Dotplot visualizing identified differentially accessible peaks (DAPs) near canonical cell marker genes across 11 cell types annotated using integrated snATAC-seq and scRNA-seq (Fig 2C) datasets. Such cell type DAPs are significantly more accessible (p_val_adj < 0.05) in one cell type compared to the average of all other cell types (see “Methods”). The nearest genes of the 12 DAPs on the x-axis from left to right were: Monocyte markers *CSF1R* and *CD14*, DCs marker: *FLT3*, B cell markers: *PAX5* and *CD19*, T cell marker *CD3E*, CD4posab marker *CD4*, CD8abPOSab marker *CD8A* and *CD8B*, NK marker *PRF1* and *KLRK1*, GD marker *TRDC*. The size of the dot represents the fraction of cells having chromatin accessibility at the DAP for each cell type. The larger dot indicates a higher percentage of nuclei with accessible region in that cell type. The color of the dot denotes the average chromatin accessibility level across all nuclei within a cell type (red is high). The genomic coordinates of the DAP genes on x axis are listed in Supplementary Table 6. The full list of cell type DAPs is described in Table 1 and Supplementary Table 7 celltype.DAP.summary. B: Dotplot visualizing gene expression of 12 marker DEGs across 11 cell types in scRNA-seq dataset. These 12 marker genes and their order on x-axis are the same as that of Fig 3A. C: Visualization of the genomic regions near the monocyte marker gene *CSF1R* described in Fig 3A. The genomic coordinate of the DAP shown in the shaded region is 151125625-151130033 on chromosome 2. The gene track and longest transcript of CSF1R is shown at the bottom of the panel. D: Visualization of different transcript of *CSF1R* created by Ensembl 102. Vertical arrows demonstrate that the Transcription Start Site (TSS) of *CSF1R*-201 and *CSF1R*-202 overlapped with the DAP described in Figure 3C.

**Table 1.** *Cicero*-based predictions of regulatory element networks acting to regulate Differentially expressed genes in specific cell types.

|  | #<br>DEG* | #<br>DAP* | #DAP<br>within<br>promoter | #CCAN with<br>promoter DAP<br>hub | #DEG with<br>promoter DAP | #CCAN associated<br>with DEG with<br>promoter DAP hub |
| --- | --- | --- | --- | --- | --- | --- |
| Monocytes | 864 | 3590 | 653 | 47 | 97 | 14 |
| B | 308 | 1333 | 237 | 104 | 35 | 33 |
| CD8abPOSab | 273 | 424 | 83 | 39 | 12 | 13 |
| CD4posab | 197 | 587 | 73 | 33 | 12 | 8 |
| CD2negGD | 141 | 574 | 123 | 0 | 9 | 0 |
| cDCs | 507 | 1731 | 262 | 132 | 28 | 26 |
| ASC | 593 | 2109 | 443 | 222 | 46 | 42 |
| NK | 242 | 1114 | 235 | 122 | 28 | 28 |
| CD2posGD | 158 | 676 | 206 | 127 | 11 | 11 |
| pDCs | 771 | 3472 | 542 | 228 | 68 | 69 |
| Total | 4054 | 15610 | 2857 | 1054 | 346 | 244 |

\*The statistical criteria are $\text{avg\_log2FC} > 0.25$ and $\text{p\_val\_adj} < 0.05$

We also identified a cis-element region covering TSS of *PAX5* that was broadly accessible, with highest accessibility in B cells (4^th^ column of Fig 3A). This element may regulate the expression of *PAX5* specifically in B cells, as PAX5 expression was noted in all B cell clusters in scRNA-seq dataset (Fig 3B). Likewise, a cis-element region including TSS of *CD4* that was differentially accessible in annotated CD4posab cells was identified (Fig 3A) and it might account for the RNA expression pattern of *CD4* in CD4posab cells (Fig 3B). Unsurprisingly, this DAP near *CD4* was also accessible in pDCs since *CD4* is also expressed in that cell type (Fig 3B). Intriguingly, this DAP is also accessible in CD8abPOSab cells (which do not express *CD4*) (albeit less so compared to CD4POSab cells), which might be due to *CD4* expression not being solely controlled by this DAP near *CD4* and there might be some other features that regulates the expression of *CD4*.

Identified cell type DAPs were then utilized to validate the integrated cell type annotation described in Fig 2B via two approaches: First, the nearest gene of cell type DAPs were extracted and labeled with corresponding human orthologous nomenclature. The gene ontology (GO) enrichment analysis was conducted using the human genes as input. The enriched GO terms lines up with the biological function of the matching cell type (Supplementary Fig 16-20). For example, the enriched terms with highest number of genes near B cell DAPs were “immune response-regulating signaling pathway” and the “enriched B cell activation” which align with the principal roles of B cells in the adaptive humoral immune system.

### 2.5 Cell type specific transcription factor activity

To detect the TFs whose binding motif were enriched in the cell type specific cis-elements detected by snATAC-seq, and thus potentially control the cell’s biological functionality, transcription factor binding motif (TFBM) enrichment analysis was performed using the cell type DAP genomic sequences as input to the HOMER package (see “Method” section) (Supplementary Table 8 TFBM.celltype.known.result.summary). Results for the top 20 enriched TFBM for each cell type are shown, clustered by their enrichment pattern across (x-axis) cell types and across transcription factor motifs (y-axis) (Fig 4). Overall, 69 unique TFs whose binding motif was enriched in cell type DAPs were identified. The gene activity of only about 32% of identified TFs were detected in respective annotated cell types in the scRNA-seq dataset. But expression of 74% of the TFs were detected in the matching or comparable cell type in bulk PBMC RNA-seq datawhich might result from the different capture efficiencies between scRNA-seq and bulk RNA seq (Herrera-Uribe et al., 2021). TFs in the same TF family were clustered together (Fig 4). It is interesting that the clustering of cell types by TFBM enrichment was fairly consistent with clustering shown in Fig 2B determined through chromatin accessibility patterns. While the TFBM enrichment pattern of the mixed CD8abPOSabT_NK cell cluster was not similar to NK cell nor of CD8abPOSab T cell clusters (Fig 4), this cluster did share similar binding motif enrichment patterns to both NK cells (Figure 4, from Elk1 through Zfp281) and CD8abPOSab cells (Figure 4, from Fli1 through Sp1). The similar motif enrichment landscape observed between CD8abPOSabT_NK, myeloid cells, and B lineage cells from AP-1 through ELF5 indicated the regulatory complexity of the CD8abPOSabT_NK cluster (which is likely a mixed population) and thus was not explored in the downstream analysis.

**Figure 4.**
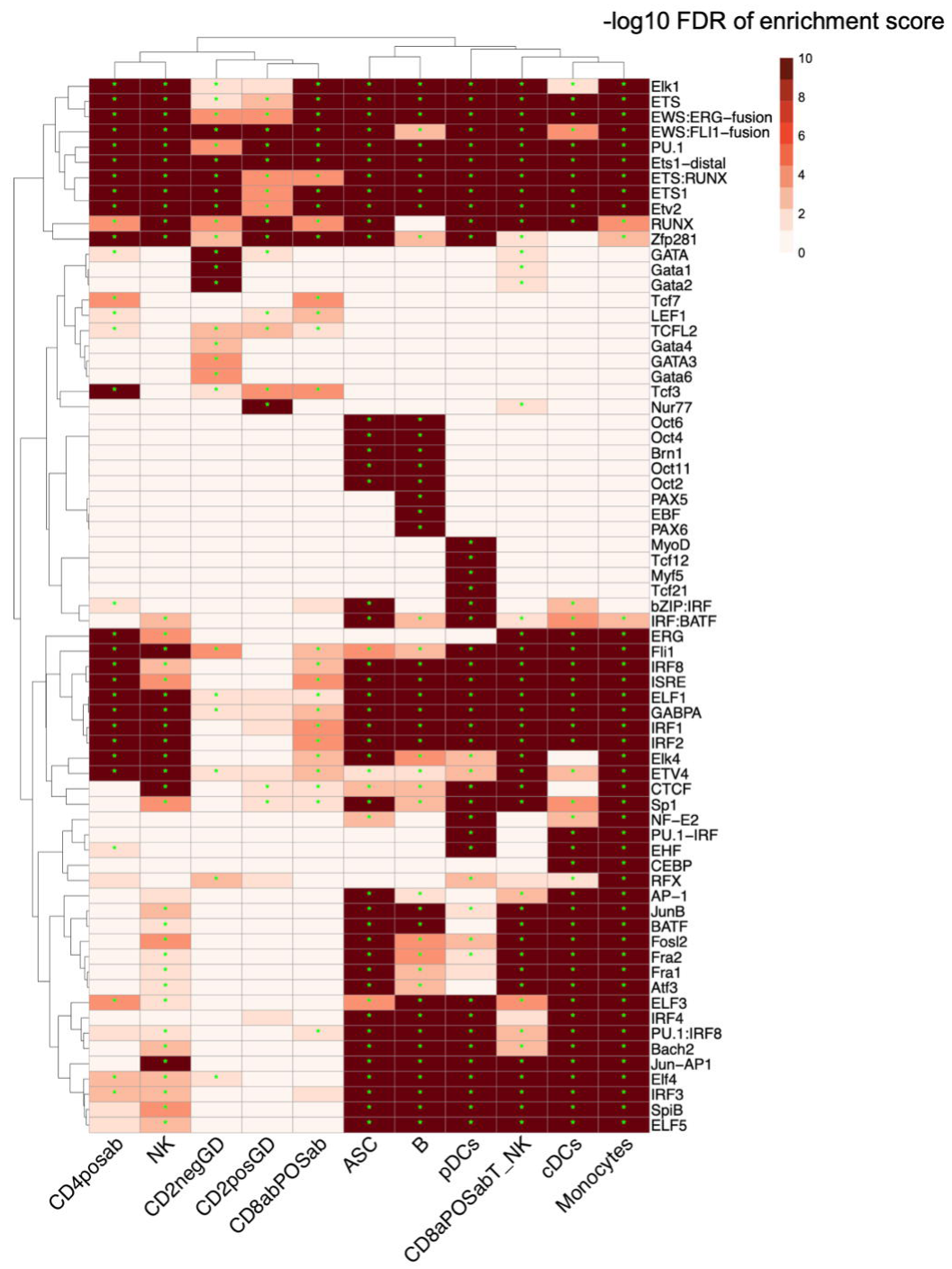
Transcription factor binding motif (TFBM) analysis of the cell type differentially accessible peaks (DAPs) predicts TF regulating these cell type networks. A: Heatmap visualizing binding motif enrichment level for top 20 TFs in each of the 11 major PBMC types. The color of the square denotes the value of -log10 of multiple test adjusted q value with Benjamini multiple testing correction. The darker color, the smaller q value and the more statistically significant. * denotes that the binding motif of the TF were statistically enriched (q < 0.05) in the corresponding cell type. The TFs and cell types were both clustered by similarity of pattern using Euclidean distance. Cell types were annotated as described in Fig 2C: Monocytes, B cells, CD8abPOSab (CD8αβ+ αβ T cells), CD4+ αβ T cells (CD4posab), CD2-γδ T-cells (CD2negGD), conventional dendritic cell (cDCs), antibody secreting cells (ASC), CD8αβ+ αβ T/NK cells (CD8abPOSabT_NK), NK, CD2+ γδ T-cells (CD2posGD), plasmacytoid dendritic cells (pDCs).

We identified both general and cell type specific TF patterns of cell type TFBM enrichment of the TFs. Several TFs had detectable enrichment of their motifs in cell types with no detectable RNA expression in the scRNA-seq dataset, like *PAX6*, indicating scRNA-seq may not be sensitive enough to detect their expression, or that the TFBM enrichment observed is unrelated to gene regulation. Unsurprisingly, the binding motif of TFs playing a crucial role in multiple immune cell types or lineages, like *PU.1* (also known as *SPI1*), *ETS*, *ETS1*, and *ETV2* were ubiquitously enriched in DAP for all cell types. We also identified a set of cell type specific TFs. The binding motif of *PAX5*, *PAX6* and *EBF* were only enriched in B cells which is compatible with the fact that *PAX5* is regarded as a B cell marker and *PAX6* has a similar binding motif to that of *PAX5*. We also predicted several TF with enriched motifs in specific cell types that have few to no reports describing them as regulators of gene expression in the immune cell type motif enrichment was observed. These included *TCF21* which has not been reported in pDCs, and *Spi-B* and *TCF12* which was predicted as candidate regulators in pDCs development (Nagasawa et al., 2008). The binding motif of *Nur77* (*NR4A1*) was most enriched in CD2posGD cells though it was also enriched in mixed CD8aPOSabT_NK cluster. The binding motif of several *GATA* family TFs (*GATA, GATA1, GATA2, GATA3, GATA4, GATA6*) were most highly enriched in CD2negGD cells. *TCF21* and *TCF12* had enrichment of binding motifs in pDCs DAP, which has also not been reported as expressed in or regulating genes specifically in pDCs. In addition, we found that *PU.1* had motif enrichment in myeloid cell DAPs (pDCs, cDCs and monocyte) through three different TF complexes (*PU.1, PU.1:IRF8*, and *PU.1-IRF*).

### 2.6 Cell type specific chromatin interactions

To predict the potential regulatory regions of DEGs (Supplementary Table 9) and predict the regulatory cis-element interactions of the TFs described in Figure 4 at specific DEGs, cis-co-accessibility networks (CCAN) analysis was performed using Cicero (Pliner et al., 2018). A CCAN is defined as a module of genomic regions that are statistically co-accessible with one another in the same cell type. To maximize the ability to link TFs to DEGs in this dataset, CCANs were predicted for each DEG with a TSS overlapping an open chromatin region that was a DAP in the matching cell type. Each CCAN has the following characteristics: 1) The “hub” peak of a CCAN overlaps with the TSS of a gene which was a DEG in the matching cell type; 2) all remaining peaks were assigned to the same CCAN as the “hub” peak if the peak has a co-accessibility score with the “hub” peak of at least 0.05 and was no more than 250,000 bp 5’ or 3’ to the gene TSS. Across 11 cell types, we identified 244 such CCANs in total (Table 1), and the total number of peaks in these CCANs ranged from 3-49. The full list of genomic regions in each predicted significant CCAN for each cell type can be found at FigShare link https://figshare.com/articles/journal_contribution/pig_PBMC_snATAC_CCAN_files_celltype_D EG_bed_files_zip/24762189 (DOI:10.6084/m9.figshare.24762189). As examples of CCAN with highest average co-accessibility score with the center peak in each cell type, we visualized CCANs associated with *POU2AF1* in B cells, *CST7* in NK cells, *MEF2C* in cDCs cells, CD5 in CD4posab cells (Fig 5), *FLNB* in ASC, *FSCN1* in CD2posGD, *ARL4C* in CD8abPOSab, *S100A8* in Monocytes and *CXorf21* in pDCs (Supplementary Fig 21).

**Figure 5.**
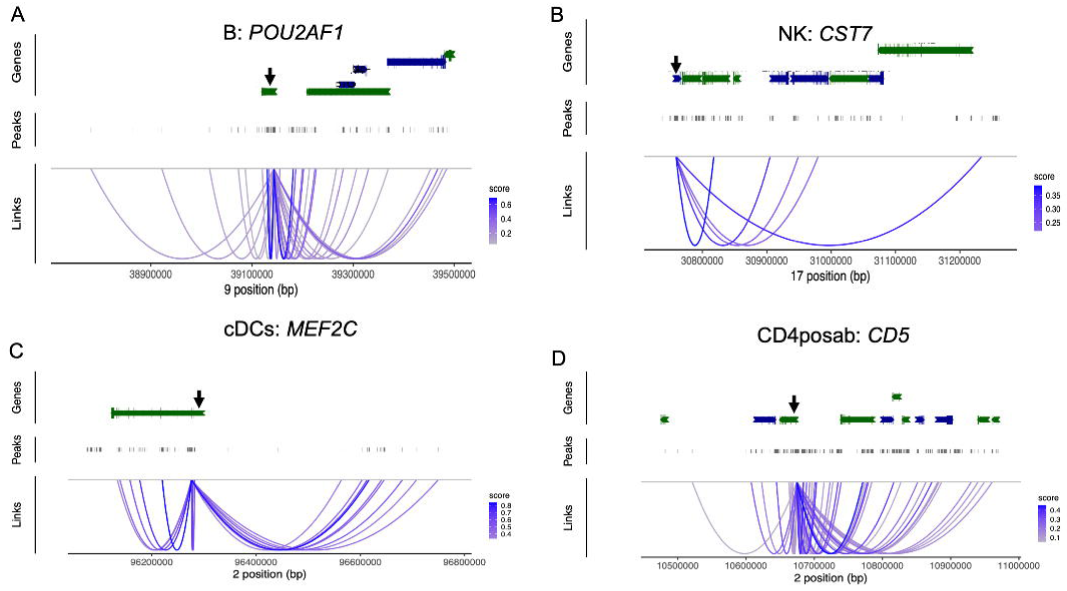
Cis-co-accessibility network (CCAN) architecture at indicated differentially expressed gene in specific peripheral immune cell types. Visualization of CCANs associated with DEGs in four different peripheral immune cell types. The center of each CCAN overlaps with the TSS of a DEG in respective cell type. Each purple line denotes that the peaks at either end of the line has a co-accessibility score greater than 0.05. Genes neighboring DEGs associated with each CCAN were not shown for the sake of clarity. A: CCAN at *POU2AF1* in B cells. The “hub” peak (chr9-39139969-39146482) of this CCAN was a DAP overlapping TSS of *POU2AF1* in annotated B cells. All peaks identified in snATAC dataset in this region are shown, but only the 24 peaks correlated with the hub peak with a co-accessibility score > 0.05 are included in the CCAN (purple). B: CCAN at *CST7* in NK cells. The “hub” peak (chr17-30755868-30762233) of this CCAN is a CD2posGD cell DAP overlapping TSS of *CST7*. There were five peaks correlated with the hub peak with a co-accessibility score > 0.05; C: CCAN at *MEF2C* in cDCs cells. The “hub” peak (chr2-96274161-96278261) of this CCAN is a cDCs cell DAP overlapping TSS of *MEF2C*. There were 19 peaks correlated with the hub peak with a co-accessibility score > 0.05; D: CCAN at *CD5* in CD4posab cells. The “hub” peak (chr2-10671164-10677736) of this CCAN is a CD4posab cell DAP overlapping TSS of *CD5*. There were 16 peaks correlated with the hub peak with a co-accessibility score > 0.05;

### 2.7 Regulators involved in cell type specific chromatin interactions

Since the TSS hub peak is co-accessible with the peaks in the rest of the CCAN, the CCAN predicts regulatory regions potentially interacting with the accessible promoter to regulate differential expression of the DEG through binding regulatory proteins (Muto et al., 2021). To predict such potential regulatory TF for DEGs, TFBM enrichment analysis was performed, using the combined regions from each CCAN as input to HOMER. The binding motif of 70 TFs (41 unique TFs) was found enriched in one or more CCANs. These motifs were associated with 45 DEG (43 unique genes) in 8 of 11 annotated PBMC cell types. Only a few of these 41 TFs were detected in scRNAs-seq dataset while gene expression of 80% of these TFs were detected in corresponding or most comparable cell type in bulk RNA-seq of sorted porcine immune cells. These differences result from the fact that bulk RNA-seq has a deeper sequencing depth(Herrera-Uribe et al., 2021). Some TFs (*ZNF519,GFY, ISRE, Fra1, Fra2, GFY, SpiB and GRE*) were not detected in bulk RNA-seq, which can be the result of the following factors: 1) Their expression levels were too low to be detected by bulk RNA-seq, 2) Since we used vertebrate motif sets to perform TFBM analysis, these TFs do not necessarily have to be expressed in porcine immune cells, 3) It could be that other expressed TFs, who are in the same family of these undetected TFs, function as real regulators since they have similar binding motif.

The results are illustrated across these CCANs through sorting by cell type and clustering by patterns of enrichment of TF motifs (Fig 6). Unsurprisingly, *CTCF* and *BORIS* (a CTCF-Like Protein) were in the same cluster and their binding motif was enriched in the CCANs of multiple genes. *IRF1* can directly bind the IFN-stimulated response element (*ISRE*) to control expression of IFN-stimulated gene regarding *IFN-I* and *IFN-II* (Michalska et al., 2018). This might explain why *IRF2*, which is also in the *IRF* family having conserved binding domain, and *ISRE* are assigned into the same cluster. In addition, our results demonstrate the value of CCANs to identify putative regulators of DEG and verify TF-related genes previously predicted in Fig 4. For example, *POU2AF1* is a transcriptional coactivator in complex with either *OCT1* or *OCT2* whose binding motif were enriched in global B cell DAPs in Fig 4. Moreover, the enriched binding motif of CTCF in the CCAN of *CST7* in CD2posGD could have contributed to the enriched binding motif of CTCF in CD2posGD DAPs in Figure 4.

**Figure 6.**
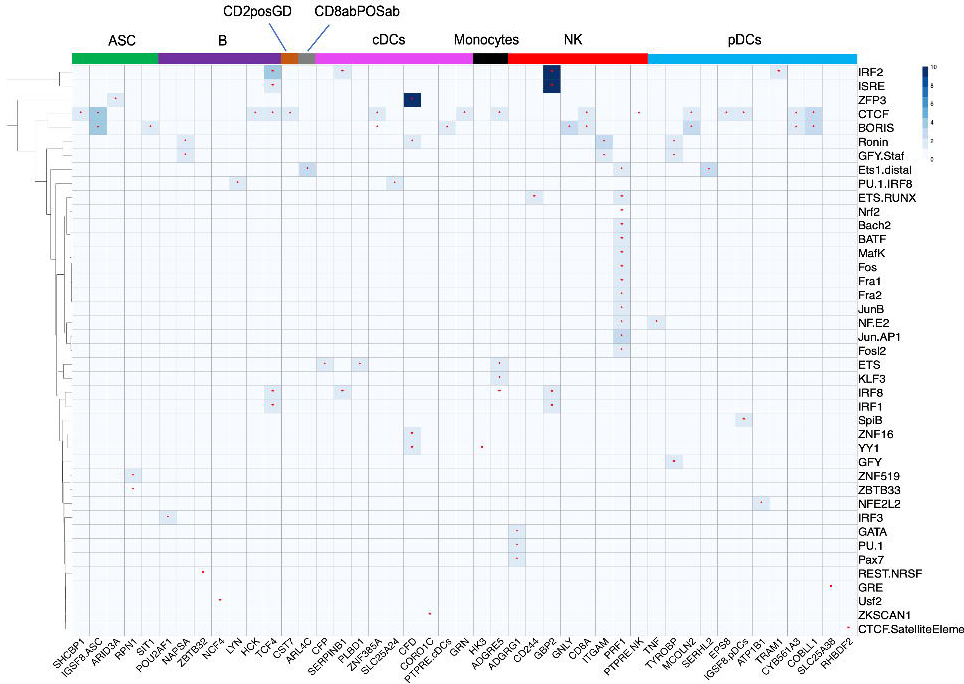
Transcription factor binding motif (TFBM) analysis on CCANs peaks identified potential regulator TFs for specific DEGs. Potential transcription factors regulating cell type specific CCANs were identified through evaluation of transcription factor binding motif analysis of cis-co-accessibility network analysis. A: Heatmap visualizing the enrichment level of all TFs whose known binding motif(s) were enriched (q <= 0.1) in at least one CCAN associated with a DEG having a DAP overlapping with its TSS. The column denotes the DEG which is the hub of the CCAN, and the cell type for which the hub gene is differentially expressed is shown. The row denotes the TFs whose binding motif are enriched across all peaks of a CCAN. The color of the cell denotes the value of -log10 of q value for enrichment. The darker color, the smaller q value and the more statistically significant. * denotes that the binding motif of the TF are statistically enriched (q < 0.1) in the peaks of the CCAN associated with corresponding DEG. The TFs is clustered using Euclidean distance.

Overall, there are 1-3 TF binding motifs enriched in each CCAN. Interestingly, exception to this observation is the binding motif of 13 TFs enriched in the CCAN associated with *PRF1* in NK cells. *PRF1* is highly expressed in NK cells and encodes a central protein (perforin) for NK cell function; thus, a highly active CCAN at the *PRF1* promoter is not surprising. One of these, *NRF2*, is known to regulate *PRF1*(Jessen et al., 2020). On the other hand, several of these TFs are sub-units of *AP-1*, a well-known general transcription factor: Fos gene family members (*FOS, FOSL2, Fra1(FOSL1)* and *Fra2(FOSL2*)) can encode protein dimerizing with proteins in Jun family (*JunB* and *Jun*) to form the *AP-1* transcription factor complex. In addition, *MAFK* or other small *MAF* proteins can bind to the same motif as *NF-E2*. Thus, this unusually large number of different TFBM enriched in the *PRF1* CCAN may be explained due to these functional overlaps for an *AP-1*-regulated gene. Interestingly, there is potential antagonistic interactions among enriched TF at *PRF1*; *BACH2* regulates transcription (activation or repression) via *MAFK*, but *BACH2* can inhibit *AP-1* proteins in blood (Lesniewski et al., 2006).

To compare our predicted TF-target gene pairs with other studies, we explored the target genes of the 41 unique TFs in the *ENCODE* Transcription Factor Targets dataset (TFTD), which was created using ChIP-seq (ENCODE, Project Consortium et al. (2004); Myers et al., 2011; et al., 2016). 22 of the 41 unique TFs shown in Fig 6 and their predicted target genes have been reported in ENCODE TFTD. Notably, for these 22 TFs, 57% (30 out of 53) predicted TF-target gene pairs described in Fig 6 were highly consistent with the *ENCODE* TFTD result (Supplementary Table 10). For example, the binding motif of *IRF3* was enriched in the CCAN peaks of *POU2AF1*, which was identified as a target of *IRF3* reported in *ENCODE* TFTD. The binding motif of *CTCF* was enriched in CCANs of 15 DEGs (14 unique DEGs) and all of these predicted target genes were concordant with those reported in the *ENCODE* TFTD. Likewise, *ZKSCAN1* was predicted to regulate *CORO1C* in cDCs and *MAFK* to regulate *PRF1* in NK cells; these relationships were also reported in the *ENCODE* TFTD. Additionally, we also found some predicted regulatory relationships were similar to what has been reported in *ENCODE* TFTD. These include the result that *ETS* family TFs are predicted to bind to *ADGRE5* is analogous to the relationship of *ETS1* and *ADGRE5* reported in *ENCODE* TFTD (Supplementary Table 10).

Further, we also have some novel findings beyond *ENCODE* TFTD. For example, TF *BATF* and its associated gene *PRF1* in NK cells were not reported in *ENCODE* TFTD. *IGSF8* is a DEG having a promoter DAP and predicted CCANs in both ASC and pDCs. Interestingly, the binding motif of both CTCF and *BORIS* are enriched in the CCAN of *IGSF8* in ASC, while the binding motif of *SpiB* is enriched in the CCAN of *IGSF8* in pDCs. Similar *IGSF8* expression level in ASC and pDCs (Supplementary Fig 13C), while enrichment of different TF in the same target gene might elucidate different regulatory mechanism governing the expression of *IGSF8* in different immune cell types through a different regulatory network. Notably, *SPIB* is predicted to be a target gene of *CTCF* in ENCODE TFTD. On the contrary, *PTPRE*, a DEG having a promoter DAP and predicted CCANs in both cDC and NK, might be regulated via similar pathways in these two cell types since the biological function of *BORIS* and *CTCF* is similar.

## 3. Discussion

A detailed functional annotation of the porcine genome will greatly improve our understanding of porcine gene regulation and network biology, as well as accelerate genetic improvement of important traits such as disease resilience. While new epigenetic data across adult tissues has provided initial chromatin state maps (Kern et al., 2021; Pan et al. 2022), there is limited information on the regulatory regions in porcine immune cells (Foissac et al., 2019; Herrera-Uribe et al., 2020). To identify such regulatory elements, we profiled the first chromatin accessibility landscape of freshly isolated porcine PBMC at single cell resolution. We demonstrated that this landscape of accessible regions at known marker genes could be explored to annotate cell type without the use of gene expression data. Integration with scRNAseq data was effective to both verify such annotations and to improve some ambiguities. Identifying regions more accessible in specific cell types was then exploited to predict TF that may bind such regulatory elements to control cell type expression. Correlation of accessibility among open chromatin regions were then used to predict both cis-co-accessibility networks (CCANs) at specific genes, as well as predict the TF controlling expression of these genes. These results were validated with ATAC-seq data from bulk-sorted PBMC populations and are consistent with many reports on specific gene regulatory factor networks.

### 3.1 Open chromatin regions detected with single nuclei ATACseq methods can be used to identify and annotate specific immune cell types in porcine peripheral blood

A deep collection of high-quality open chromatin regions was identified and approximately 22% of these accessible regions were within 3kb from the TSS of an annotated gene (Fig 1A). This fraction was relatively low compared to that reported for human (in kidney; Muto et al., 2021). It might originate from the fact that the pig genome is not annotated as well as that of human or that this is the characteristic of immune cells compared to tissues; however, we detected much higher TSS enrichment scores [average was 17.99 (Supplementary Fig 1B-E)] than that of human PBMC snATAC-seq whose average is 12.55(Wu et al., 2022). Our two replicates showed high similarity and were integrated into a dataset of 17,207 nuclei and grouped into 35 clusters. By utilizing the DNA accessibility patterns of putative cis regulatory elements (gene activity) as a proxy for gene expression, this “gene activity” measure at several known gene markers for major cell types was used to annotate the 35 clusters. Gene activity for canonical gene markers was estimated with two methods: assigning all peak data (< 2,000 bp from TSS) to the closest gene, and by calculating the DAP for each cluster and using the specific DAP mapping proximal to the marker gene for estimating gene activity. We observed that the gene activity scores created from all nearby accessibility data were less definitive than the pattern(s) for DAPs at the canonical marker genes (Fig 1C-D, Supplementary Fig 6, Supplementary Table 4), which may demonstrate the most important regulatory elements for cell type expression may be TSS-proximal DAP. Using the TSS-proximal DAP approach, we classified cell type by inspecting DAP patterns near all markers whose scRNAseq patterns of expression were used as cell type markers in PBMC (Herrera-Uribe et al., 2020). Monocytes, B, ASC, DCs, T, CD4posab, CD8abPOSabT_NK, NK, GD and unknown cells were determined sequentially. Consequently, the 35 clusters were grouped into 13 cell types (Fig 2B).

Comparing the chromatin accessibility pattern in Fig 2A and the corresponding gene expression pattern in Herrera-Uribe et al., 2021 two general DAP-cell type expression patterns were observed:

1. The DAP and the cell type where this DAP is open were consistent with the expression of the nearest gene in Herrera-Uribe et al., 2020. For instance, some B clusters (0, 8) and ASC (27) at *CD86* DAP2, ASC (27) at *CD19* DAP, cDCs (26) at *CSF1R* and *CD86* DAPs, DC(26,29) at *TCF4* DAP1, monocytes (5,7,9) at *FLT3* DAP, DC(26,29), ASC (27) and CD8abPOSab (13, 19, 24, 30, 32) and CD8aPOSabT_NK (25, 33) at *SLA-DRB1* DAPs, monocytes, B cells, ASC, CD8abPOSab, NK and CD8aPOSabT_NK at *XBP1* DAP, CD8abPOSab at *PRF1* DAP, pDCs (29) at *CD4* DAP2.
2. Accessible chromatin patterns had no nearby gene with matching scRNAseq gene expression reported (Herrera-Uribe et al., 2020. For example, ASC (27) demonstrates DNA openness at *PAX5* DAP without revealing *PAX5* expression in Herrera-Uribe et al., 2020. Similar patterns were found in CD8abPOSab (24, 30) at CD86 DAP2, CD2posGD (22) at *XBP1* DAP, NK (10), CD8abPOSab (30) and B (8) at *TRDC* DAP, pDCs (29) at *CD8B* DAP2. The chromatin accessibility at the DAP near *CD4* in cluster 21,23, and 30 in Fig 2A might demonstrate the complexity of the gene expression and there are potentially multiple regulatory regions controlling the expression of this gene. These patterns might originate from the complicity of regulative mechanism in biology, the heterogeneity of a known porcine immune cell type and the variability of the sensitivity of snATAC-seq and scRNA-seq. Multiple regulatory elements and TFs can contribute to the regulation for the same gene coordinately and thus, the DNA openness at one cis-element near a gene might not necessarily pair with the gene expression in a cell type since they could miss an essential element to activate the gene expression compared to the cell types where the gene is expressed. Notably, we did not detect a peak overlapping the *CD2* gene that would distinguish CD2posGD and CD2negGD directly, although there are a few DAPs whose nearest gene is *CD2* (6,893bp or more distant, Supplementary Table 4). Because of this distance we did not use these peaks to predict *CD2* status or for cell type determination. We divided DC (26, 29) into sub types via the chromatin accessibility at *SLA-DRB1*, *CD8A, PRF1 and KLRK1* cis-elements since they were highly expressed in CD2posGD but not CD2negGD cells, though they were not defined as CD2posGD markers (Herrera-Uribe et al., 2021).

### 3.2 Chromatin accessibility pattern annotation verified and improved through integration with scRNAseq data

To further explore and validate these proposed annotations, the snATAC-seq data was integrated with previously published scRNAseq data, and a high level of validation was observed. We showed that the chromatin accessibility pattern of cis regulatory elements near the cell type markers used in Herrera-Uribe et al., 2020, was highly similar to the pattern of the expression level of matching DEG (Fig 2B-2C, Supplementary Table 5). Our results revealed the snATAC-seq has similar power to scRNA-seq in terms of cell type annotation, although there were a small number of inconsistencies. But its ambitious to define the cell type definitely using only using the peaks, since there are usually multiple open chromatin regions near one gene and now the prior knowledge about which particular region is more informative than others in terms of one marker is limited. Considering the cell type assignment outcomes are nearly uniform in Fig 2B-C, the cell types of snATAC-seq predicted using scRNA were grouped into the same cluster as matching sorted porcine PBMCs in bulk ATAC-seq of in Supplementary Fig 3B-3C and the predicted cell type by integrating with scRNA in Fig 2C is more definitive compared to that in Fig2B, we used the predicted cell types using scRNA-seq to conduct further analysis.

After studying the characteristic of cell type DAP near a TSS of a gene, we found that our dataset predicted the prospective TSS specifically used in the matching porcine immune cell types. Among DAPs including TSS of characterized cell markers in Herrera-Uribe et al., 2020, we predicted that 2 DAPs overlay the cell type specific TSS candidates for *CSF1R* and *CD8A* (Figure 3C-D and Supplementary Fig15).

### 3.3 Porcine PBMC cell type regulatory elements were enriched for transcription factors known to control immune cell differentiation and function

A characteristic of cell type regulatory elements is that they can also be used to identify putative regulatory factors through TFBM enrichment analysis. This can be especially useful to complement regulatory network analysis using scRNAseq alone, since scRNA-seq is sparse and insensitive for detecting lowly expressed TFs. Thus, we defined the TFs that lead to the cell type specific biological functionality and that function ubiquitously across multiple pig immune cells by recognizing the TFs whose binding motif are enriched in cell type DAPs (Fig. 4). The binding motif of *PU.1 (SPI1), ETS, ETS1, ETV2, Elk1, EWS, RUNX and Zfp281* were predicted to be comprehensively enriched in diverse cell type DAPs. Besides, *PU.1* was predicted as regulators in myeloid cells (pDCs, cDCs and monocytes) via 3 schemes: *PU.1*,*PU.1:IRF8*, and *PU.1-IRF*. We observed several TFs (*POU5F1*, *POU2F3*, *POU2F2* and *POU3F3*) in the *POU* domain family whose binding motif are enriched in ASC and B cell DAPs. The observed specificity of *Oct2* (*POU2F2*) in B cells is consistent with what has been previously reported (Küppers, 2021). We also have identified several known cell-specific TF. *Nur77*, encoded by the *NR4A1* gene and whose binding motif was enriched in CD2posGD cell DAPs in F, was reported to be expressed in pig Treg cells and has been recently shown to mediate T cell differentiation even during immunosuppression by calcineurin inhibitors (Sekiya et al., 2022). Our finding that binding motif of a couple of *GATA* family TFs including *GATA3* are most enriched in CD2negGD cells agrees with the observation that *GATA3* is highly expressed in pig CD2negGD cells compared to other GD cells (Rodríguez-Gómez et al., 2019; Gu et al., 2022), as well as the *GATA3* gene expression pattern reported in Herrera-Uribe et al., 2021 (Supplementary Fig 13B). At the same time, some novel regulators were predicted in this TFBM analysis. The predicted regulative function of *Spi-B* and *TCF12* in pDCs DAPs supported a sparsely studied interaction (Nagasawa et al., 2008).

### 3.4 Cis-acting regulatory networks and transcription factor-target gene relationships predicted from correlating chromatin accessibility of regulatory elements

The prediction for the involvement of a TF in regulating genes through DAP for each cell type above did not attempt to connect a specific TF and its target gene(s). To define the regulatory networks with a higher resolution, the TFs were linked to DE target genes in each cell type via CCAN generation and identifying the regulatory network for such genes. The summary of number of CCAN associated with a DEG in each cell type was provided in Table 1. The fact that no such CCAN in CD2negGD might be due to the fact that CD2negGD has the least number of DEG with a promoter DAP, making it less possible to construct enough peak connections to assemble a CCAN associated with these DEG. Since the peaks are co-accessible with the DAP overlapping with a TSS of a DEG and are mostly within a window of size of 500,000 bp, motivated by the fact that scientist found the peaks near a gene are highly consistent with the regulatory enhancer region of identified using Chip-seq (Muto et al., 2021), we assumed that the promoter or enhancer region of that DEG can be covered in the genomic regions in the CCAN. Driven by the aim of exploring the prospective regulator for DEG, we performed TFBM analysis for each CCAN having a hub peak overlapping with a TSS of DEG. As a consequence, our outcomes summarized in Fig 6 indicate that CCANs are a powerful means to recognize regulator candidates of DEG and potentially refine the TF-target genes described in Fig 4. Additionally, our predicted TFs and their related DEG were highly consistent with or similar to regulatory relationships predicted in the ENCODE TFTD. This cross-species verification provided evidence that our predicted relationship between a regulator and its target gene may often be correct. Nevertheless, we also pinpoint some either new or porcine-specific TF-target gene pairs. Our results demonstrate the great power and sensitivity of snATAC-seq to elucidate the chromatin accessibility landscape of pig immune cells, determine the known cell types based on the DNA open element pattern, predict the regulators for each cell type, and create the first resource of TF and possible target genes, including the matching possible binding sites, in different unstimulated porcine immune cell types.

## 4. Limitations

We recognize several limitations that constrained our power and likely accuracy. Firstly, we have only two biological replicates of PBMC from a single timepoint and pig breed. However, open chromatin regions in our replicates were very consistent, and by using un-manipulated PBMC we avoided potential changes to cell transcriptomes that can occur with the extensive sorting that would be required to collect large numbers of rare cells, such as ASC or DC. Our ability to exclude potential breed biases reflected in the results from adult Yorkshire pigs is limited, but this first dataset provides a foundation that can be expanded. In addition, our published scRNA and snATAC were profiled from different samples. Profiling gene expression and chromatin accessibility from the same cells could be helpful to avoid the integration of these two ‘omic’ datasets. However, the integration produced a combined cell type annotation was highly consistent between omics methods.

## 5. Conclusions

The genome-wide catalog of regulatory elements in this snATAC-seq dataset, including the cell type DAPs and the regulatory elements in the CCAN at a DEG are important resources to improve genome-wide genetic variation analyses. One example use of these data is filtering of variants associated with important phenotypes such as disease resilience and resistance in pig populations, as a majority of disease- and trait-associated noncoding Genome-wide association study (GWAS) variants are localized in this type of genomic regions (Maurano et al., 2012). The predicted TF-target gene network is also a highly useful resource for future characterization of the regulatory elements controlling porcine immune cell identity for immunology and biomedical

## 6. MATERIALS AND METHODS

### 6.1 PBMC sample collection, nuclei isolation and snATAC-seq using 10x Chromium

Four PBMC samples were isolated from 2 healthy 6-month-old FAANG founder male Yorkshire pigs using standard techniques(Herrera-Uribe et al., 2021). PBMC nuclei were isolated by following DEMONSTRATED PROTOCOL: Nuclei Isolation for Single Cell ATAC Sequencing (10x Genomics) with an adjustment: The concentration of nuclei suspension, which were stained by Ethidium homodimer-1, was measured and determined using Countess II FL Automated Cell Counter. Then 4 libraries from two batches were constructed as described in Chromium Next GEM Single Cell ATAC Reagent Kits v1.1 (10x Genomics) and sequenced via Illumina Novaseq 6000 sequencing runs at DNA facility at Iowa State University.

### 6.2 Demultiplexing and generation of single-cell accessibility counts

Porcine genome reference and gff3 file were downloaded from ensembl 102 and used to generate the config to create a reference package using cellranger-atac mkref function of Cell Ranger ATAC (V.1.2.0). Then the base call files (BCLs) were demultiplexed using cellranger-atac function to produce the FASTQ files. For each library, the single cell accessibility counts matrix was generated using the customized reference package by cellranger-atac count command.

### 6.3 Nuclei doublet detection

To remove the doublet resulting from droplet that contains two cells, nuclei doublet detection using ArchR (1.0.1) was performed on R 4.1.1. The geneAnnotation was created using createGeneAnnotation function of ArchR with customized org and TxDb packages for Sus scrofa as input. ArrowFile for each of the dataset was constructed by running ArchR function createArrowFiles with the fragment files generated by Cell Ranger ATAC, geneAnnotation and genomeAnnotation scrofa genome Sscrofa 11.1 as input with default parameter. Inferred doublet score for each cell was added to each of the Arrow file using addDoubletScores function with default parameter. An ArchRProject was created by running ArchRProject function with generated arrow files as input. Then, 1466 detected nuclei doublets were filtered out with filterDoublets function with default filterRatio. The cell barcodes of non-doublets were pulled out for downstream analysis.

### 6.4 Quality control, snATAC-seq datasets integration, and clustering

The detected peaks using cellranger-atac in two datasets from animal 6798 and 6800 were merged using reduce function of GenomicRanges (1.42.0), respectively (Lawrence et al., 2013). The merged 6798 peaks having overlaps with merged 6800 peaks were merged with merged 6800 using subsetByOverlaps of GenomicRanges. The peaks with a width >= 10000 bp or <=20 bp were filtered out from the merged peaks of two animals to generate a set of high-quality unified peaks. The fragments detected was counted in this new set of peaks using FeatureMatrix command of Signac (1.4.0), a ChromatinAssay was created by CreateChromatinAssay of Signac with min.features = 1000 and a Seurat (4.0.5) object was created using CreateSeuratObject for each of the dataset. The cells predicted to be doublets by ArchR were removed every Seurat object. Low-quality cells were removed from 4 Seurat objects (nucleosome_signal < 4, TSS.enrichment > 2, nCount_peaks > 2000, nCount_peaks < 30000) before term frequency inverse document frequency (RunTFIDF) normalization. 4 Seurat objects were merged and visualized by RunUMAP with dims = 2:30 as input. A set of integration anchors were defined by FindIntegrationAnchors and used as input to integrate 4 Seurat objects by running IntegrateEmbeddings using 1:30 dimensions of merged Seurat object. The integrated snATAC-seq Seurat object was normalized and its most variable features were identified by RunTFIDF and FindTopFeatures, respectively. The correlation between sequencing depth and every reduced dimension component was checked by DepthCor. 2:30 reduced dimensions of the integrated Seurat object were used to define 35 clusters by running “FindClusters” with a resolution = 2.4 using shared nearest neighbor (SNN) clustering algorithm. A bar plot was created to visualize the percent of cells in each cluster from each dataset. The DAP and the number of DAP in all pairwise clusters were summarized in Supplementary Table 1-2.

### 6.5 Cell type annotation for clusters using snATAC-seq

Regulatory regions potentially controlling cluster-specific gene expression were identified by measuring DAP for each cluster using FindAllMarkers of Seurat with min.pct = 0.2, logfc.threshold = 0.25, only.pos = TRUE. The list of DAPs for each cluster was provided in Supplementary Table 3. Predicted gene activity profiles were created using two ways: 1) by counting the Tn5 transposase cutting sites in fragments of nearby genes (<2000 bp from TSS). Particularly, the overall estimated gene activity of the gene markers used in Herrera-Uribe et al., 2021 were used to decide the cell types for clusters. This was used to roughly narrow down the possible clusters for a cell type (Supplementary Fig 6-11). 2) by counting the Tn5 transposase cutting sites at a cluster DAP whose nearest gene is one of the gene markers used in Herrera-Uribe et al., 2021 (Supplementary Table 4)(Herrera-Uribe et al., 2021). This criterion was used to determine the cell types more precisely in Fig 2A-B. The example comparison of the predicted gene activities using these two methods are provided in Supplementary Fig 12. The principles reflected in Fig 1C, Supplementary Fig 12 and Supplementary Table 4 to annotate the cell types are described as below.

A. The first cell type determined was monocyte (cluster 5, 7, and 9) by checking the chromatin accessibility at 5 DAPs near *CSF1R*, *CD14* and *CD86* (first 5 columns in Fig 1C).
B. The next cell type decided was B cells (cluster 0,3, 4, 8, 11, 18, 20) based on the chromatin openness at 2 DAPs near *PAX5* and *CD19* (6^th^-7^th^ column in Fig 1C, Supplementary Fig 12 A-B). Then cluster 27 was defined as ASC using 3 DAPs near *PRDM1* and *TCF4* (*TCF4* was highly expressed in ASC though it was not classified as an ASC marker in Herrera-Uribe et al., 2021)(8^th^-10^th^ columns in Fig 1C).
C. DC clusters (26 and 29) were decided based on a DAP near *FLT3* and was further interpretated as cDCs (26) based on 3 DAPs near *SLA-DRB1* and pDCs (29), with an elevated Tn5 cutting sites in 4 DAPs at *XBP1*, *IRF8*, *IRF8* and *CD4*.
D. Chromatin accessibility at a DAP neighboring *CD3E*, identified T cell clusters (1, 2, 6, 12, 13, 14, 16, 17, 19, 21, 22, 23, 24, 25, 28, 30, 32, 33, 34). Subsequently, clusters 2, 6 and 12 were characterized as CD4posab since the cells are largely accessible at *CD4* DAP. Detection of chromatin openness at 3 DAPs near *CD8B* and *CD8A* enabled the definition of CD8abPOSab (13, 19, 23, 24, 30, 32). Afterwards, cluster 10 was characterized as NK due to the lack of chromatin accessibility at *CD3E* DAP and the openness at *PRF1* DAP and *KLRK1* DAP. CD8abPOSabT_NK (25 and 33) was determined since the cells demonstrate the chromatin openness at *CD3E*, *CD8A*, *PRF1* and *KLRK1*. *TRDC* gene activity was investigated to define GD cells (1, 14, 16, 28 and 22). Furthermore, the presence/absence of DNA accessibility at *SLA-DRB1*, *CD8A*, *PRF1* and *KLRK1* DAPs were used to classify CD2posGD (22) and CD2negGD (1, 14, 16, 28) since these genes are highly expressed in CD2posGD Herrera-Uribe et al., 2021 though they were not described as CD2posGD marker. Cluster 17, 21 and 34 were grouped into a particular subtype of T cells due to the co-accessibility of chromatin near markers of various cell types.
E. Cluster 15 and 31 was determined as unknown cell type since it has elevated estimated gene activity for *PAX5*, *XBP1*, *CD3E*, *PRF1* and *TRDC*.

### 6.6 Cell type annotation for snATAC clusters by integration with scRNA dataset

To further annotate the cell types, the cell types were predicted for each cell by integrating snATAC-seq with our published PBMC scRNA-seq data(Herrera-Uribe et al., 2021). A set of anchors were detected by running FindTransferAnchors having estimated gene activity of snATAC-seq as query and scRNA-seq data as reference with reduction = ‘cca’. The most possible cell type labels predicted for each of the cell in snATAC-seq dataset were transferred to snATAC-seq by TransferData with the new reduction of integrated snATAC-seq as weight.reduction and dims = 2:30. The cells in snATAC-seq with a low prediction.score.max <= 0.5 were excluded from our Seurat object.

### 6.7 Cell type DAP identification

Based on the predicted cell type labels by integrating with scRNA-seq, the genomic regions differentially accessible in one cell type compared to the average of all other cell types were detected by running FindAllMarkers function of Seurat with min.pct = 0.1, only.pos = TRUE, logfc.threshold = 0.25 and p_val_adj < 0.05. The list of such DAPs for each cell type was provided in Supplementary Table 7 celltype.DAP.summary.

### 6.8 Comparison with bulk ATAC-seq

We identified shared peaks between scATAC and ATAC-seq of bulk sorted porcine PBMC populations using bedtools intersect with reciprocal overlap > 25% between peaks(Quinlan and Hall, 2010). Read counts in common peaks were obtained using featureCounts(Liao et al., 2014), and principal component analysis was applied using base R software to visualize clustering of scATAc-derived and bulk sorted PBMC populations. Enrichment and corresponding significance of cell type DAPs within DAPs from bulk sorted porcine PBMC populations were calculated using hypergeometric tests in base R.

### 6.9 GO analysis of the genes close by cell type specific DAPs

The nearest gene of the cell type DAPs were found and then converted to matching human homologous via Ensembl 102. Further, GO analysis was performed for each cell type using Metascape with the corresponding human genes as input(Zhou et al., 2019). The ontology terms in which the input genes are enriched were detected using hypergeometric test and Benjamini-Hochberg p-value correction algorithm with all genes in the genome as background. Enriched terms were groups into clusters and Kappa-test score was used to capture the most representative term for each cluster. Further, the most significant terms with a Kappa score above 0.3 in each cluster were kept. The networks are all visualized via Cytoscape(Shannon et al., 2003). The GO analysis result was summarized in Supplementary Figure 16-20.

### 6.10 Transcription factor binding motif analysis of cell type DAPs

TFs associated with each cell type and might act as important regulators in each cell type. TFBM analysis was performed using HOMER with this setting “-size given -mask -mset vertebrates” to discover the TFs whose binding motif are enriched in exact size of cell type DAPs compared to the GC-content matching background peaks generated from the Sscrofa11.1 genome. The q value of a TF binding motif was calculated by taking the average q values of this TF in the corresponding cell type DAP if its binding motif is enriched in one cell type DAPs via different co-factors. The threshold of q value of TF binding motifs was set as 0.05. The known motif enrichment results for each cell type were listed in Supplementary Table 8 TFBM.celltype.known.result.summary.

### 6.11 Generation of cis-co-accessible networks using Cicero

Chromatin cis-co-accessibility analysis was performed using R package Cicero (1.8.1). Seurat object of each cell type was converted to CellDataSet format via as.cell_data_set and then used to generate input for cicero by make_cicero_cds function. The co-accessibility score for all peak pairs on each chromosome of Sscrofa 11.1 was calculated using the generated CellDataSet object by running run_cicero. All pairwise peaks are filtered following these criteria: 1) At least one of the pairwise peaks are a DAP in the matching cell type, 2) The co accessibility score of the pairwise peaks are greater than 0.05. The peaks meet the criteria above were grouped into co-accessible networks using generate_ccans with default setting. The constructed CCANs were further refined as below: 1) The center peak of the CCAN overlaps with a TSS of a DEG in the matching cell type. 2) All peaks in the CCAN were assigned with the same CCAN number.

### 6.12 Prediction of the regulator for DEG in each cell type

TFBM was conducted using HOMER with following setting “-size given -mask -mset vertebrates -N 300” to predict the TFs whose binding motif are enriched in peaks of each CCAN described above compared to the GC-content matching background peaks generated from Sscrofa11.1 genome. The threshold of q value of TF was set as 0.1.

## Data availability Statement

Raw sequencing data from snATAC-seq are available through the European Nucleotide Archive (project: PRJEB68307 (SAMEA8050928) at https://www.ebi.ac.uk/ena/browser/view/PRJEB68307 and PRJEB68308 (SAMEA8050929) at https://www.ebi.ac.uk/ena/browser/view/PRJEB68308). The scripts used for this study can be found at https://github.com/pengxin2019/snATAC_PBMC_2023_Tuggle.

## Ethics Statement

The animal study was reviewed and approved by USDA-ARS-NADC Animal Care and Use Committee.

## Author Contributions

CT and CL conceptualized and supervised research. JH-U, KB, and CL collected and cryopreserved PBMC samples. PX-Y performed nuclei isolations, supervised the sequencing and analyzed the snATAC-seq dataset. PX-Y, RC, CL, and CT interpreted the data and drafted the manuscript. RC provided the bulk ATAC-seq data from flow-sorted cell populations. LD assisted PX-Y with early bioinformatics analyses. All authors contributed to the writing of the materials and methods, edited the manuscript, and approved the final version.

## Acknowledgements

This work is supported by National Institute of Food and Agriculture (NIFA) Project 2018-67015-2701 and USDA-ARS CRIS 5030-31320-004-00D. We thank the DNA facility at the Iowa State University for technical support and sequencing platforms used in this study. In addition, we thank the NADC animal care staff for their efforts. We also thank Dr. Jayne Wiarda for explaining the scRNA-seq dataset and giving suggestions on data analysis. The data analysis of this work was performed on high performance computing Nova cluster of Iowa State University.

